# Shifts in diversification influence rates of song evolution in honeyeaters

**DOI:** 10.1101/2023.06.27.546781

**Authors:** Eleanor M. Hay, Steven L. Chown, Matthew D. McGee

## Abstract

Birdsongs are acoustic signals that can play a role in sexual selection. Despite the established role of birdsong in selection and reproductive isolation at microevolutionary scales, the macroevolutionary relationships between song evolution and clade diversification remain largely unexplored. We test the hypothesis that shifts in diversification influence rates of song evolution in honeyeaters, a diverse clade of songbirds restricted to Australasia. Using a song dataset for 163 honeyeater species, we identify lineage-specific shifts in diversification, and test the influence of these shifts on song evolution with state-dependent, relaxed, multivariate Brownian motion models. We also explore the underlying patterns of song evolution and phylogenetic signal of song. Shifts to lower rates of diversification are associated with lower rates of song evolution. This suggests the two factors are correlated, but is contrary to expectations that elevated rates of diversification will be associated with elevated rates of song evolution. We also found that song follows a punctuated mode of evolution, implying that birdsong has an important role in speciation events, although this signal likely erodes over time and is subject to phylogenetic constraints.

## INTRODUCTION

Determining what factors cause species diversification is a central goal of many evolutionary studies. The influence of morphological traits (Cooney & Thomas, 2021; Rabosky et al., 2013; Simões et al., 2020) and environmental factors such as ecological opportunity (Dumont et al., 2012; Mahler et al., 2010), on rates of speciation and diversification have been a major focus, but other factors including sexual signals, can also play a role (Maan & Seehausen, 2011; Ritchie, 2007; Wagner et al., 2012). Divergence and development of reproductive isolation are key initial phases in speciation events (Coyne & Orr, 2004; Hernández-Hernández et al., 2021; Jarvis, 2021). Sexual signals frequently emerge as the most divergent traits between closely related species and often experience strong selection (Ritchie, 2007; Servedio & Boughman, 2017). The evolution and divergence of sexual signals can outpace ecological divergence (Arnegard et al., 2010). Consequently, it has been hypothesized that rapid changes in sexual signals could drive species diversification (Maan & Seehausen, 2011; Servedio & Boughman, 2017).

Vocalizations may act as sexual signals that influence speciation. Acoustic signals are used by a wide range of organisms and are complex behavioral traits that enable individuals of the same species to identify one another (Laidre & Johnstone, 2013). Acoustic signals have broad-scale functions in species recognition and fine-scale functions in mate choice, thus potentially influencing speciation (Catchpole & Slater, 2003; Wilkins et al., 2013). Divergence of acoustic signals has played a role in the early stages of speciation in a wide range of organisms including cicadas (Marshall et al., 2008), crickets (Gray & Cade, 2000; Tinghitella et al., 2018; Zuk et al., 2006), frogs (Boul et al., 2007), and birds (Grant & Grant, 2006). In the context of reproductive character displacement, however, the strength of selection on signals depends on range overlap, with sexual signals expected to be under greater divergence when species are sympatric rather than allopatric (Maan & Seehausen, 2011). Therefore, at macroevolutionary scales (i.e., clade or family-level), the strength of selection on acoustic signals is expected to vary between species and among clades. Whether the processes of divergence, assortative mating, and selection of acoustic traits, which occur at the microevolutionary scale, can have macroevolutionary consequences for clade diversification, remains unclear.

Birdsong is one of the best-known acoustic signals and is an important sexual signal. Male birds sing to attract females. Consequently, song can act as an important premating isolation barrier and influence speciation (Price, 2008; Tobias et al., 2020). Indeed, song evolution has long thought to be a factor that could explain the diversity of passerine birds (Baptista & Trail, 1992; Fitzpatrick, 1988). The role of birdsong in speciation and reproductive isolation has been studied at fine scales: empirical evidence supports the theoretical expectations. Divergence of song has contributed to reproductive isolation in warblers (Irwin et al., 2001) and is thought to have influenced the early stages of peripatric speciation in antbirds in Amazonia (Seddon & Tobias, 2007). Additionally, long-term studies on Darwin’s finches have demonstrated that sexual selection on differences in vocalizations, resulting from changes in beak morphology, have impacted mate choice and contributed to reproductive isolation and speciation (Grant & Grant, 2006; Podos, 2001; Podos & Nowicki, 2004). Although acoustic signals are commonly hypothesized to influence diversification (Wilkins et al., 2013), most studies have focused on population or sister species comparisons. Empirical studies testing the influence of signal divergence on speciation at broader scales remain scarce (Mason et al., 2017). Where the relationship between birdsong and speciation has been investigated at macroevolutionary scales, bursts in rates of speciation coincide with bursts in song evolution, however, no significant correlations have been uncovered at the family-level (Mason et al., 2017).

Indeed, the relationship between diversification and birdsong has proven difficult to disentangle, largely due to song itself being a complex trait, which in birds is influenced by developmental mode and constrained by factors such as beak morphology, body size, and the environment (Derryberry et al., 2018; Friedman et al., 2019; Friis et al., 2022). The majority of macroevolutionary studies on birdsong have therefore focused on the underlying drivers of song evolution. Like most sexual signals, acoustic signals are expected to diverge between close relatives, especially if ranges overlap, potentially leading to a lack of phylogenetic signal in the trait (Pfennig & Pfennig, 2010). However, song frequency is understood to be phylogenetically conserved (Mikula et al., 2021), largely due to constraints imposed by morphological traits, especially body size (Derryberry et al., 2018; Pearse et al., 2018). Additionally, the developmental mode of song is also thought to influence song evolution and could complicate its relationship with speciation. Birdsong ranges from learned to innate, and learning is understood to be a form of phenotypic plasticity that is influenced by cultural transmission and can impact evolutionary outcomes (Pfennig et al., 2010). Learned songs are typically associated with higher rates of evolution (Mason et al., 2017) and divergence (Lachlan & Servedio, 2004) than innate songs, although the evidence is mixed (Freeman et al., 2017). Such complexity in the evolution of acoustic traits therefore provides a challenge for diversification approaches and can result in disparity when investigations are conducted over varying phylogenetic and spatial scales.

To overcome such limitations, we test the relationship between birdsong evolution and diversification using a monophyletic clade restricted to a particular region, leveraging the power that concentrating on a diverse, single clade can bring (Marquet et al., 2004). Thus, we focus on honeyeaters (Aves: Meliphagidae), a phenotypically diverse clade of birds (192 species; Gill et al., 2020) that range throughout Australasia and are known for their distinctive vocalizations (Higgins et al., 2008). Song in honeyeaters is learned and varies greatly among species, with some species exhibiting simple and monotonal songs, whereas others have complex and variable songs (Fig. 1). This variation in vocalizations, combined with the high diversity and endemism of honeyeaters, makes the group ideal for studying the evolution of vocalizations and their potential role in species diversification.

**Figure 1.**
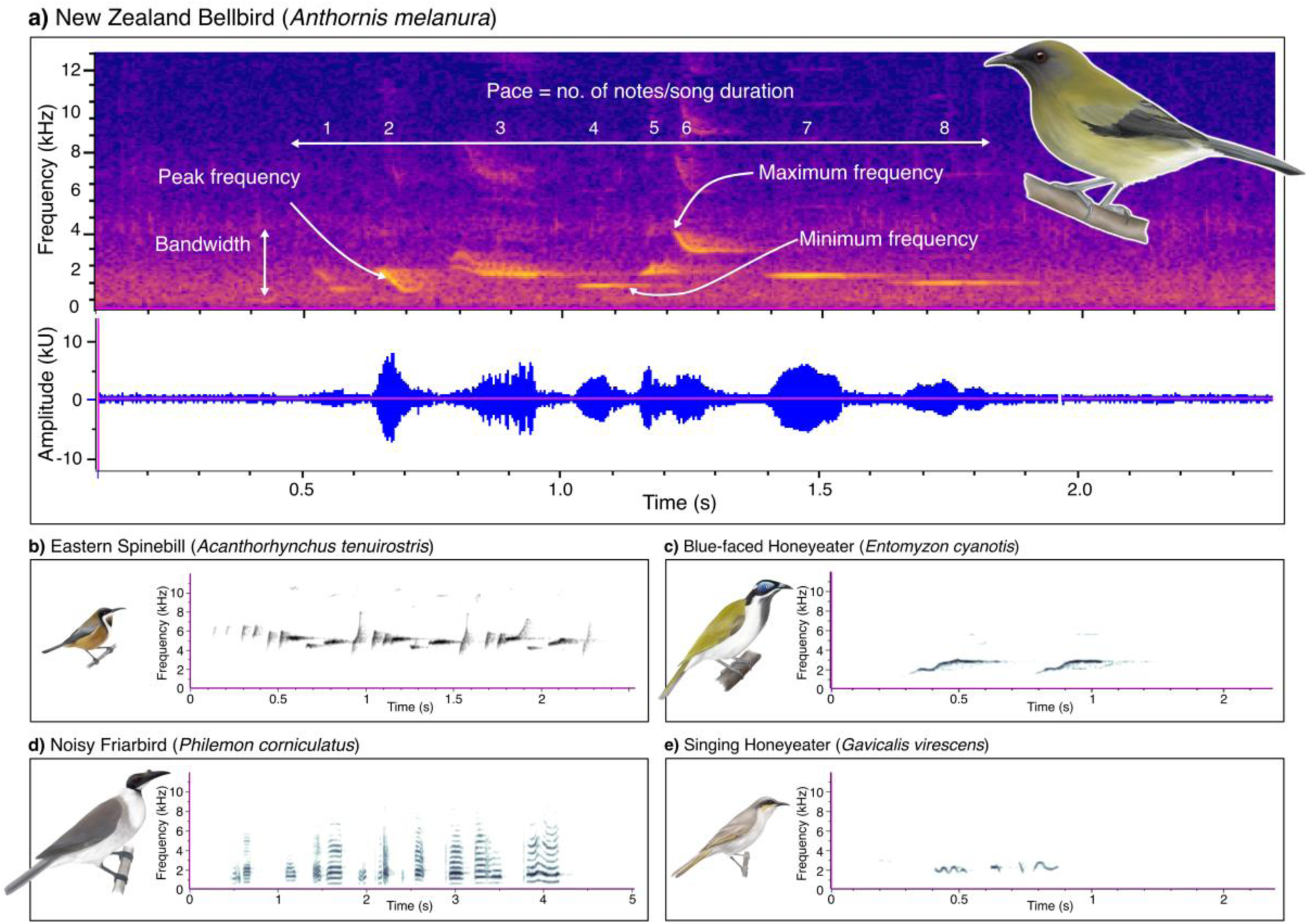
Song spectrograms displaying frequency (kHz) over time (s) for five species of honeyeater: a) New Zealand Bellbird (*Anthornis melanura*; xeno-canto accession number = XC153005), b) Eastern Spinebill (*Acanthorhynchus tenuirostris;* xeno-canto accession number = XC490800), c) Blue-faced Honeyeater (*Entomyzon cyanotis*; xeno-canto accession number = XC439288), d) Noisy Friarbird (*Philemon corniculatus*; xeno-canto accession number = XC287056), and e) Singing Honeyeater (*Gavicalis virescens*; xeno-canto accession number = XC334251). The five song variables considered in this study are indicated with annotations in a): peak frequency represents the frequency of maximum amplitude across the entire song, maximum frequency is the highest pitched note in song, minimum frequency is the lowest pitched note in song, bandwidth is the difference between maximum and minimum frequency across the song, and pace is the number of notes over song duration.

Specifically, we use honeyeaters as a model clade to explore the relationship between diversification and birdsong evolution. Using a similar framework to past studies that have investigated the relationship between song evolution and diversification across bird clades (Mason et al., 2017), we first test for shifts in honeyeater diversification and then assess if shifts in diversification are correlated with differential rates of song evolution. We predict that if birdsong plays a role in diversification, then shifts in diversification will influence rates of song evolution: species with elevated rates of diversification should have elevated rates of song evolution. To further disentangle the patterns of song evolution, we also compare fit of different evolutionary models to assess patterns of song evolution. We assess whether there is evidence of changes in song occurring at speciation events, which would be expected if song has a role in speciation. Lastly, we test for phylogenetic signal of song. By doing so, we can assess the theoretical expectation that if bird song is a sexually selected trait and undergoing selection from reproductive character displacement, then there will be a lack of phylogenetic signal.

## METHODS

### Song and phylogenetic data

We used a previously collated dataset of male honeyeater song variables (Hay et al., 2024). Containing data for 163 honeyeater species, the song dataset considers five aspects of song (Fig. 1; Table S1) that have been explored in past macroevolutionary birdsong studies (e.g., Derryberry et al., 2018; Mason & Burns, 2015; Mason et al., 2017): peak frequency (frequency in which the most sound energy was produced), maximum frequency (maximum frequency of the highest pitched note in song), minimum frequency (minimum frequency of the lowest pitched note in song), song bandwidth (maximum frequency - minimum frequency), and song pace (number of notes/song duration). Song variables were log_10_-transformed prior to statistical analysis to meet parametric assumptions of normality and homogeneity of variance, and because logarithmic scales of sound frequency correspond to how birds perceive and modulate sound (Cardoso, 2013).

To estimate diversification rates and model trait evolution, we used a honeyeater phylogeny from Hay et al. (2022). This phylogeny contains all 192 species of honeyeater (Gill et al., 2020), and was constructed using a combination of nuclear genes, mitochondrial genes, and topological constraints from previous phylogenomic studies.

### RevBayes models

A range of phylogenetic comparative methods implemented through the program RevBayes (Höhna et al., 2016) were used to test the relationship between diversification and birdsong evolution. RevBayes enables Bayesian inference for phylogenetic comparative methods, and uses a probabilistic graphical model framework, which is a powerful generic framework for specifying and analyzing statistical models (Höhna et al., 2014; Höhna et al., 2016).

First, to explore patterns of diversification in honeyeaters, we estimated branch-specific speciation and extinction rates (Höhna et al. 2019) using RevBayes (version 1.2.4; Höhna et al., 2016). This model uses birth-death processes, in which diversification rates vary among branches, to estimate lineage-specific rates of diversification. Because speciation and extinction models are subject to bias when species are missing (Mynard et al., 2023), we estimated branch-specific speciation and extinction rates using the full honeyeater phylogeny containing 192 species (Hay et al., 2022). Estimates of the number of diversification rate shifts are sensitive to priors (Höhna et al. 2019), therefore we repeated this analysis using several priors on the number of rate shifts (3, 5, 8, and 10 rate shifts). The MCMC sampler was set for 4 runs of 2,500 generations and a burn-in of 10%. Model convergence was determined using Tracer v1.7.1 (Rambaut et al., 2018) by ensuring ESS were above 200. Output from RevBayes models was visualized in R (R Core Team, 2013) using the package ‘RevGadgets’ (Tribble et al., 2021). The number of shifts in diversification rate and the lineages associated with shifts were identified by extracting the ‘num_shifts’ variable in R. Priors had little influence on the number of diversification rate shifts and species associated with shifts in diversification (Fig. S1-4): we present the results from the model with 5 rate shifts.

To test the influence of shifts in diversification rates on rate of song evolution, we then used state-dependent, relaxed, multivariate Brownian motion models of trait evolution (MuSSCRat; May and Moore 2020) in RevBayes (Höhna et al., 2016). These models enabled us to test the influence of shifts in diversification identified by the previous model (a discrete character: species associated with shift in diversification vs not associated with shift in diversification) on the evolution of the five song variables (multivariate continuous characters: peak frequency, maximum frequency, minimum frequency, song bandwidth, and song pace). The MuSSCRat model is advantageous because it jointly estimates the evolutionary history of the discrete and continuous characters, while also considering background rate variation (variation in the evolution of the continuous character, not due to the discrete character of interest). This method permits rates to vary along branches and among continuous characters and reduces biases in rate estimates. For this analysis, phylogeny was trimmed to the 163 honeyeater species we have song data for using the ‘*drop.tip’* function from the ape package (Paradis et al. 2004) in R (R Core Team, 2013). The MCMC was run for 500,000 generations with a 10% burn-in. Tracer v1.7.1 (Rambaut et al., 2018) was used to visualize output and determine convergence. To evaluate the sensitivity of posterior parameter estimates, we repeated the MCMC across different priors on the number of rate shifts of the continuous characters (5, 10, and 15 shifts). The R package ‘RevGadgets’ (Tribble et al., 2021) was used to visualize the state-dependent branch rates with the ‘*plotTrace*’ function, and summary statistics were calculated using *‘summarizeTrace’* (Table S2). To further explore the underlying patterns of song evolution, we also extracted the background rates and overall Brownian motion rates of song evolution estimated from these models.

Correlations between shifts in diversification and rates of song evolution were visualized using the R package ‘phytools’ (Revell, 2012).

### Kappa models and phylogenetic signal of song

To further explore underlying patterns of song evolution and how this might be linked with diversification, we compared fit of different evolutionary models and tested for phylogenetic signal of song.

We fitted different models of trait evolution, specifically testing for trait changes at speciation events, which would be expected if song influences speciation. We used Kappa models, which transform branch lengths based on trait changes, and compared the fit of these models against Brownian motion models in which traits are assumed to evolve at a constant rate. Higher Kappa values typically indicate gradual evolution, or anagenesis, whereas Kappa values below 0.5 suggest a punctuated mode of evolution, and that character states change at speciation events. To fit these models, we used the function ‘*transformPhylo.ML*’ from the ‘MOTMOT’ package (Thomas and Freckleton, 2012) in R.

Trait values are predicted to be more similar between closely related species, however sexual signals are often expected to be the most divergent traits between species (Servedio & Boughman, 2017). Therefore, we tested how conserved song variables are in honeyeaters. The phylogenetic signal of each song variable was estimated using Pagel’s lambda (Pagel, 1999) and a likelihood ratio test using the ‘*phylosig’* function from the ‘phytools’ package (Revell, 2012) in R. Pagels lambda is a scaling parameter that ranges from 0 to 1. Using this measure, a lambda of 0 indicates no correlation between species, and a lambda of 1 indicates that the correlation of trait values between species is equal to expectations of Brownian motion (Revell et al., 2008).

## RESULTS

Branch-specific speciation and extinction models identified several patterns of honeyeater diversification. Overall, there were two diversification rate categories found for honeyeaters. Much of the tree follows a relatively constant rate of diversification; however, the model identifies several species associated with a rate shift to a lower rate of diversification (Fig. 2). Species associated with shifts in diversification are typically from the more basal linages within each clade, many of which are monotypic genera (e.g., *Glycicharea fallax*, *Certhionyx variegatus*, *Oreornis chrysogenys*, *Purnella albifrons*).

**Figure 2.**
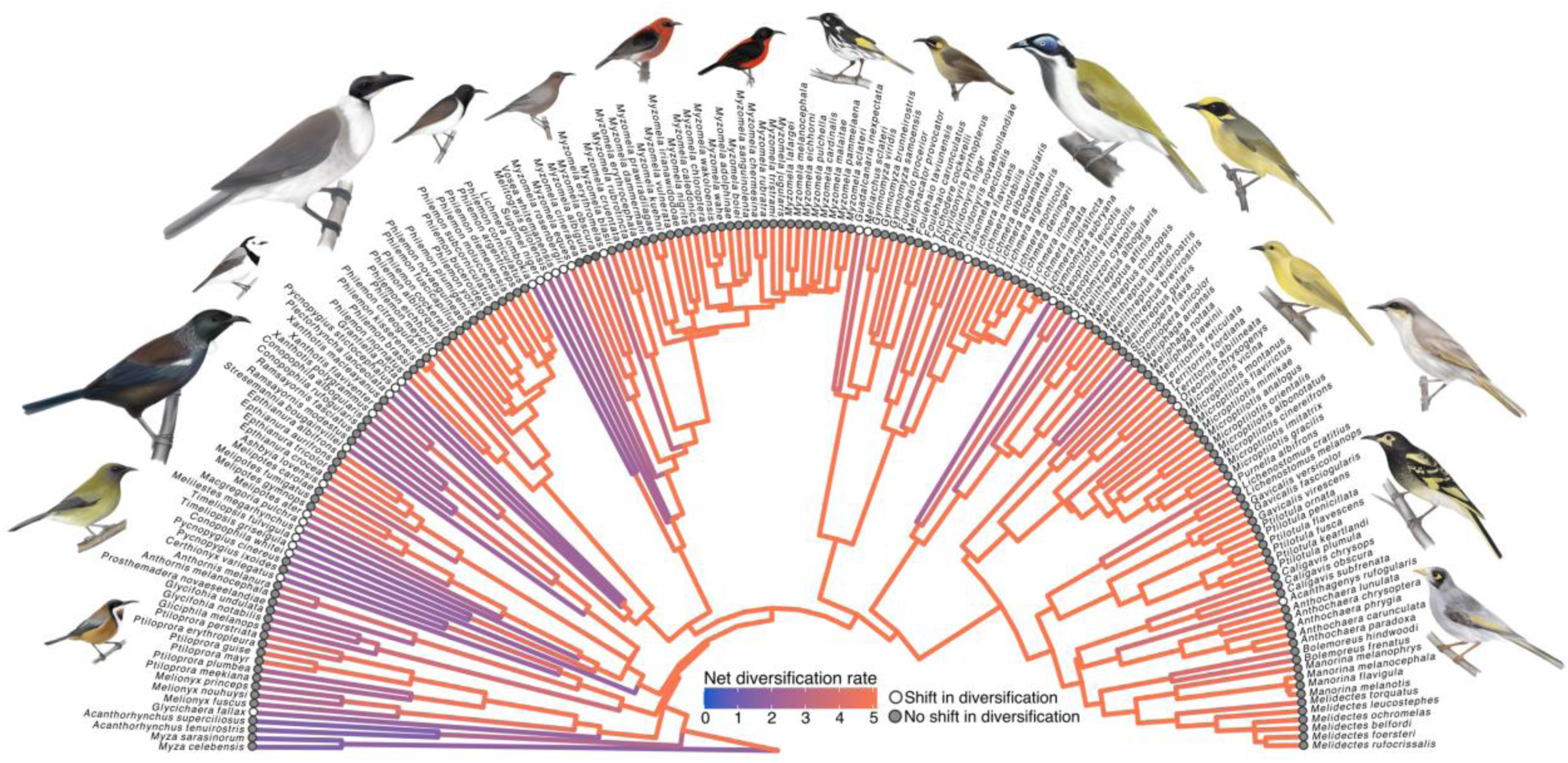
Lineage-specific net diversification rates for honeyeaters. This is modeled across the full phylogeny of 192 honeyeater species. Branches on the phylogeny are colored according to diversification rate, with the warmer colors representing high rates of diversification and cooler colors representing lower rates of diversification. Lineages associated with shifts in diversification are indicated with the dots at the tips of the phylogeny.

MuSSCRat models revealed that shifts in diversification have a significant influence on rates of honeyeater song evolution (posterior probability = 1.0). This was consistent across different priors (Fig. S5-7). The state-dependent branch rates show that shifts to lower rates of diversification are associated with lower rates of song evolution (Fig. 3a). There is no overlap of confidence intervals and over a 10-fold difference in song evolution rate estimates between these two diversification rate categories (mean rate of song evolution_diversification shift_ = 0.17, 95 % CI = [0.091, 0.31]; mean rate of song evolution_no diversification shift_ = 1.83, 95 % CI = [1.69, 1.91]; Fig. 3c).

**Figure 3.**
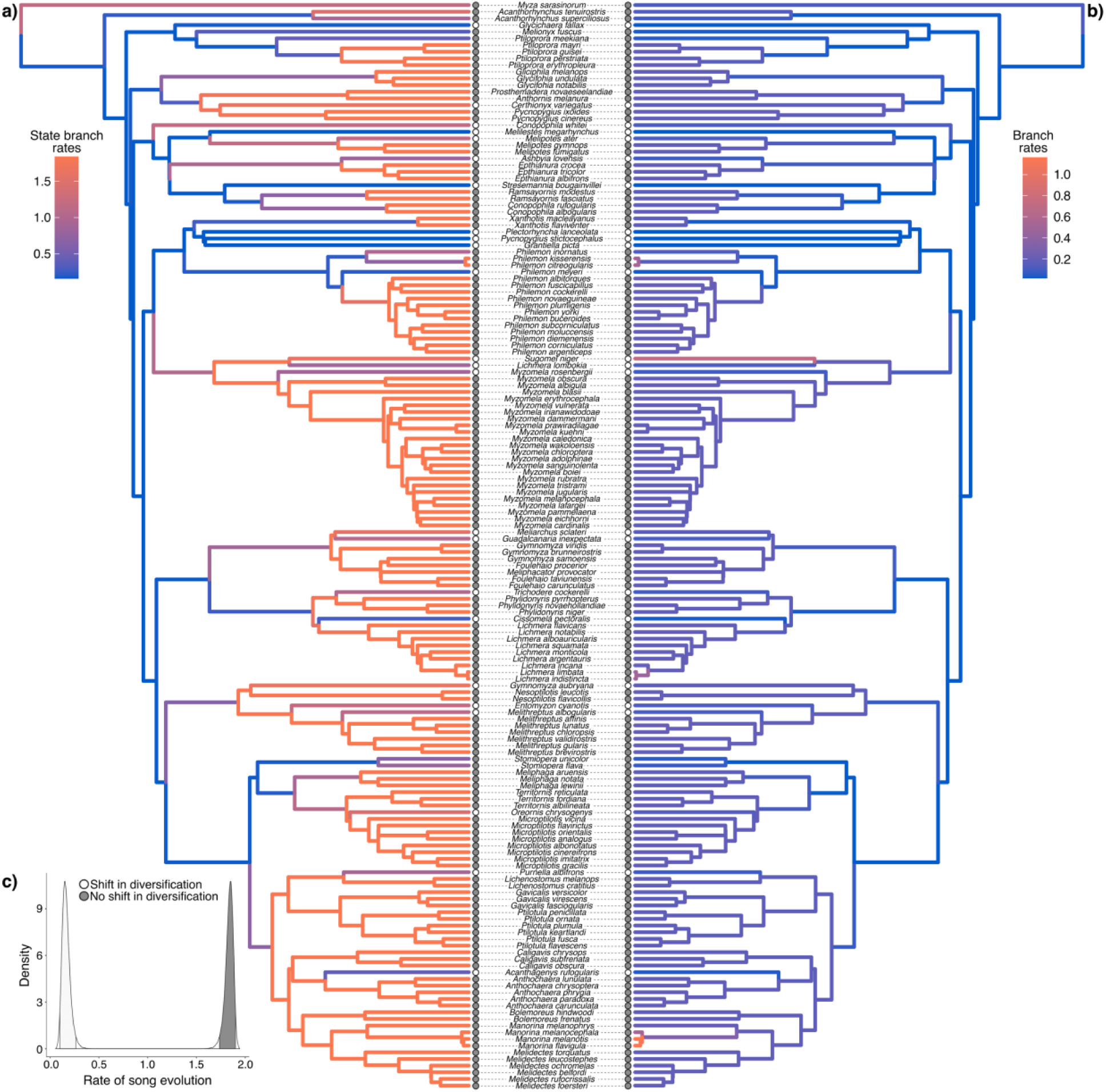
Patterns of honeyeater song evolution extracted from MuSSCRat models. Here, analysis was completed using the 163 honeyeater species we had song data for. Branches on the phylogeny on the left (a) indicate state-specific branch rates. The states tested are indicated with the dots at tip labels, these are the species that were found to have shifts in diversification. Rates of song evolution for the two states are displayed in the density plot (c). The phylogeny on the right (b) displays the overall Brownian motion rates of song evolution extracted from models.

The overall Brownian motion rates of song evolution extracted from MuSSCRat models uncover further patterns of song evolution (Fig. 3b). Most of the phylogeny exhibits a relatively constant rate of birdsong evolution, however increased rates of evolution are notable in some young sister species. For example, *Philemon kisserensis* and *Philemon citreogularis* are associated with increased rates of song evolution, as well as *Lichmera limbata* and *Lichmera indistincta,* and *Manorina melanocephala*, *Manorina melanotis*, and *Manorina flavigula* (Fig. 3b). *Sugomel niger* is also identified as having high overall rates of song evolution.

Comparison of different trait evolution models provides further insight into honeyeater song evolution. The Kappa models perform better than the Brownian motion models in all cases (Table S3). Values of Kappa extracted from models are all below 0.5, suggesting that song follows a punctuated mode of evolution, with changes in trait values being associated with cladogenesis. Maximum frequency returned the lowest Kappa statistic (0.34), followed by peak frequency (0.35), song pace (0.38), bandwidth (0.46), and minimum frequency (0.50).

Lastly, we found that the phylogenetic signal varied substantially among the five song variables. Values of Pagel’s lambda ranged from 0.31 to 0.92 (Table S4). Song bandwidth had the lowest signal (λ = 0.31), followed by maximum frequency (λ = 0.58), pace (λ = 0.66), peak frequency (λ = 0.75) and then minimum frequency (λ = 0.92).

## DISCUSSION

We found that shifts in rates of diversification for honeyeaters are correlated with significant differences in rates of song evolution. Theory suggests that if song has a role in speciation and diversification of clades then bursts of diversification should correlate with bursts in song evolution (Wilkins et al., 2013). Past studies following a similar approach to the one adopted here found relationships between bursts in song evolution and speciation across 581 species of tanagers and ovenbirds, but this was not significant when each family was considered separately (Mason et al., 2017). We found that shifts to a lower rate of diversification are significantly correlated with lower rates of song evolution, providing evidence that song evolution and diversification are associated within families.

The directionality of this finding is, however, unusual given that expectations are typically for increased rates of diversification to be associated with increased rates of song evolution if song is acting as an isolating mechanism (Baptista & Trail, 1992; Price, 2008; Tobias et al., 2020). Honeyeater species associated with shifts to lower diversification rates are comparatively older lineages to the rest of the tree and consist of many monotypic genera. Older lineages have more time to accumulate species, but they also have more time for extinction. In birds, rates of speciation tend to slow over time as niches become filled (Ricklefs 2006; Price 2008). The lower rates of diversification we observe in these older lineages have likely resulted from speciation rates decreasing and extinction remaining constant over time. It’s important to note that the directionality of our finding is partly due to the way that the analyses have been conducted: the diversification model found shifts to a lower rate of speciation, and this was correlated to lower rates of song evolution. Using this approach there was technically no way to relate shifts to higher diversification rates (since we did not find these) to higher rates of song evolution. Nonetheless, we still find two overall rate categories of diversification and song evolution in honeyeaters, in which lower rates of diversification are correlated with lower rates of song evolution and higher rates of diversification are correlated with higher rates of song evolution.

Our results do raise the intriguing question of what might cause rates of song evolution to slow down in honeyeaters. At the lineage level, complexity and diversity of honeyeater songs is known to be impacted by population density. For example, the critically endangered Regent Honeyeater (*Anthochaera phrygia*) has had the complexity of their songs decline for decades (Crates et al. 2021). Song in honeyeaters is learned and young juveniles must co-occur with other breeding adult males to learn songs (Crates et al. 2021). Regent Honeyeaters are becoming so rare that young birds are unable to learn their own songs. This has consequences for mating success of individuals since males sing to attract female birds and can cause further population declines and even extinction. One explanation for our findings may be that such processes at the population level can scale up in the extinction-speciation interaction to have the macroevolutionary consequences on clade richness we observe in our study.

Exploring underlying patterns of song evolution enabled us to further unveil the relationship between song evolution and diversification. We find evidence that song follows a punctuated rather than gradual model of evolution, suggesting that changes in honeyeater song are important for speciation events. Overall Brownian motion rates of song evolution extracted from MuSSCRat models (Fig. 3b) corroborate this punctuated mode of song evolution and show increased rates of song evolution associated with some recent speciation events, however, we do not see increased rates with older speciation events. One possible explanation is that past signal of song divergence and evolution at speciation events has eroded over time, which could be why we observe increased rates of song evolution in young sister species. In general, success in linking micro with macroevolution has been variable (Schluter 2024): past studies linking birdsong and diversification have had mixed outcomes. For example, a study of 175 taxon pairs of New World passerine birds found no association between faster rates of song evolution and higher speciation rates (Freeman et al., 2022). Schluter (2024) suggests that difficulties in linking rates of reproductive trait evolution and species diversification could be because present day trait values might not accurately represent the past. It has been proposed that initial rate of species accumulation could be strongly influenced by the rate of evolution of reproductive isolation, however, over time other ecological processes that influence clade diversity intervene (Schluter, 2024). This could provide an explanation for mixed outcomes from past studies, and why the overall Brownian motion rates of song evolution only show elevated rates in young species.

The overall Brownian motion rates of evolution highlight elevated rates of song evolution in *Sugomel niger* and three young sister species groups: The Gray Friarbird (*Philemon kisserensis*) and the Little Friarbird (*P. citreogularis*); The Indonesian Honeyeater (*Lichmera limbata*) and The Brown Honeyeater (*L. indistincta*); and the Noisy Miner (*Manorina melanocephala*), Black-eared Miner (*M. melanotis*), and Yellow-throated Miner (*M. flavigula*). Songs are thought to diverge slowly in allopatry, and only rapidly increase on secondary contact, contributing to reinforcement (Lachlan & Servedio, 2004; Servedio & Boughman, 2017). Interestingly, the *Lichmera* and *Philemon* species are not currently sympatric, but do have adjacent ranges to some extent: The Indonesian Honeyeater ranges throughout Timor and the Lesser Sunda Islands, whereas the Brown Honeyeater is found across Australia, New Guinea, and other parts of Indonesia (Higgins et al., 2022), while the little friarbird ranges on Australia and New Guinea (Higgins et al., 2020a) and the gray friarbird is restricted to small islands offshore of East Timor (Higgins et al., 2020b). In saying that, honeyeaters have high dispersal ability, and there has been shifting biomes on mainland Australia and connection and disconnection of Australia and New Guinea (Andersen et al., 2014; Hay et al., 2022; Marki et al., 2017). Furthermore, Peñalba et al. (2018) have shown that population structure and geneflow is very dynamic in the little friarbird species complex and the brown honeyeater species complex and does not adhere to geographic barriers across northern Australia and New Guinea. Thus, these species have likely had contact during the speciation process, and the increased rates of song evolution we observe in these species could have contributed to divergence. The three *Manorina* species all have overlapping ranges across Australia: The Yellow-throated Miner ranges across nearly all of Australia, except for some areas along the East Coast (Higgins et al., 2020c); The Noisy Miner ranges along a large proportion of the East side of Australia (Higgins et al., 2020d); and the Black-eared Miner has the smallest range but is found inland around the intersection between South Australia, Victoria, and New South Wales (Higgins et al., 2020e). These species arguably have the highest rates of song evolution and could imply that sympatry has contributed to divergence of song. As an important caveat to this study, many of these effects are influenced by how species are classified. For example, the Indonesian Honeyeater was previously identified as a distinct species (Gill et al. 2020) but has now been synonymized with the Brown Honeyeater (Gill et al. 2023). How one classifies species and subspecies can influence outcomes of macroevolutionary analysis, although this is likely to have little effect in this context, given the strength of the relationship we found, and that our models controlled for such potential background rate variation.

We also find that song exhibits phylogenetic signal, and that most of the interspecific diversity of song frequency and pace in honeyeaters can be explained by evolutionary history. Theory suggests that divergence is expected for sexually selected traits between closely related species (Maan & Seehausen, 2011; Servedio & Boughman, 2017). Thus, the level of phylogenetic signal uncovered for frequency and pace is contrary to such theoretical expectations. This is unsurprising, however, given other studies have found similar levels of phylogenetic signal for song traits in birds: peak frequency was found to have a lambda of 0.87 for 5085 species of passerine birds (Mikula et al., 2021), and a lambda of 0.74 for 1022 passerine species (Friis et al., 2022). Our results are consistent with these previous studies and confirm that peak frequency of song is phylogenetically conserved. Phylogenetic conservatism of song is, to some extent, driven by allometric constraints imposed by body size (Fletcher, 2004). Body size itself is a conserved trait that directly impacts the length of the vocal tract and the size of the syrinx (Bertelli & Tubaro, 2002; Suthers & Zollinger, 2008; Friis et al. 2021), and is a driver of honeyeater song evolution (Hay et al., 2024). These other underlying processes likely contribute to the difficulty in disentangling the links between song evolution and diversification.

Nonetheless, we find some evidence that song evolution has contributed to diversification within this family of birds. Understanding the influence of reproductive isolation on diversification has been recognized as an urgent question in evolutionary biology (Matute & Cooper, 2021), especially because the evolution of reproductive isolation often defines speciation (Coyne & Orr, 2004). Here, we contribute to understanding of this question by testing the hypothesis that shifts in diversification are associated with increased rates of song evolution in honeyeaters. We found that shifts to lower rates of diversification are correlated with lower rates of song evolution, and that song follows a punctuated model of evolution rather than gradual, suggesting that changes in song occur at speciation events and this is followed by periods of stasis. Ultimately, we find that song evolution and diversification are correlated in honeyeaters, and changes in song play an important role in speciation events, however this signal likely erodes over time due to underlying constraints of song evolution and other ecological processes.

## REFERENCES

Andersen, M. J., Naikatini, A., & Moyle, R. G. (2014). A molecular phylogeny of Pacific honeyeaters (Aves: Meliphagidae) reveals extensive paraphyly and an isolated Polynesian radiation. Molecular Phylogenetics and Evolution, 71, 308–315. 10.1016/j.ympev.2013.11.014

Arnegard, M. E., McIntyre, P. B., Harmon, L. J., Zelditch, M. L., Crampton, W. G., Davis, J. K., Sullivan, J. P., Lavoue, S., & Hopkins, C. D. (2010). Sexual signal evolution outpaces ecological divergence during electric fish species radiation. The American Naturalist, 176(3), 335–356. 10.1086/655221

Baptista, L. F., & Trail, P. W. (1992). The role of song in the evolution of passerine diversity. Systematic Biology, 41(2), 242–247. 10.2307/2992524

Bertelli, S., & Tubaro, P. L. (2002). Body mass and habitat correlates of song structure in a primitive group of birds. Biological Journal of the Linnean Society, 77, 423–430. 10.1046/j.1095-8312.2002.00112.x

Boul, K. E., Funk, W. C., Darst, C. R., Cannatella, D. C., & Ryan, M. J. (2007). Sexual selection drives speciation in an Amazonian frog. Proceedings of the Royal Society B: Biological Sciences, 274(1608), 399–406. 10.1098/rspb.2006.3736

Cardoso, G. C. (2013). Using frequency ratios to study vocal communication. Animal Behaviour, 85(6), 1529–1532. 10.1016/j.anbehav.2013.03.044

Catchpole, C. K., & Slater, P. J. B. (2003). Bird Song: Biological Themes and Variations. Cambridge University Press.

Cooney, C. R., & Thomas, G. H. (2021). Heterogeneous relationships between rates of speciation and body size evolution across vertebrate clades. Nature Ecology and Evolution, 5(1), 101–110. 10.1038/s41559-020-01321-y

Coyne, J. A., & Orr, H. A. (2004). Speciation. Sinauer Associates.

Crates, R., Langmore, N., Ranjard, L., Stojanovic, D., Rayner, L., Ingwersen, D., & Heinsohn, R. (2021). Loss of vocal culture and fitness costs in a critically endangered songbird. Proceedings of the Royal Society B, 288(1947), 20210225. 10.1098/rspb.2021.0225

Derryberry, E. P., Seddon, N., Derryberry, G. E., Claramunt, S., Seeholzer, G. F., Brumfield, R. T., & Tobias, J. A. (2018). Ecological drivers of song evolution in birds: disentangling the effects of habitat and morphology. Ecology and Evolution, 8(3), 1890–1905. 10.1002/ece3.3760

Dumont, E. R., Davalos, L. M., Goldberg, A., Santana, S. E., Rex, K., & Voigt, C. C. (2012). Morphological innovation, diversification and invasion of a new adaptive zone. Proceedings of the Royal Society B: Biological Sciences, 279(1734), 1797–1805. 10.1098/rspb.2011.2005

Fitzpatrick, J. W. (1988). Why so many passerine birds? A response to Raikow. Systematic Zoology, 37(1), 71–76. 10.2307/2413195

Fletcher, N. H. (2004). A simple frequency-scaling rule for animal communication. The Journal of the Acoustical Society of America, 115(5), 2334–2338. 10.1121/1.1694997

Freeman, B. G., Montgomery, G. A., & Schluter, D. (2017). Evolution and plasticity: divergence of song discrimination is faster in birds with innate song than in song learners in Neotropical passerine birds. Evolution, 71(9), 2230–2242. 10.1111/evo.13311

Freeman, B. G., Rolland, J., Montgomery, G. A., & Schluter, D. (2022). Faster evolution of a premating reproductive barrier is not associated with faster speciation rates in New World passerine birds. Proceedings of the Royal Society B: Biological Sciences, 289(1966), 20211514. 10.1098/rspb.2021.1514

Friedman, N. R., Miller, E. T., Ball, J. R., Kasuga, H., Remes, V., & Economo, E. P. (2019). Evolution of a multifunctional trait: shared effects of foraging ecology and thermoregulation on beak morphology, with consequences for song evolution. Proceedings of the Royal Society B: Biological Sciences, 286(1917), 20192474. 10.1098/rspb.2019.2474

Friis, J. I., Sabino, J., Santos, P., Dabelsteen, T., & Cardoso, G. C. (2021). The allometry of sound frequency bandwidth in songbirds. The American Naturalist, 197(5), 607–614. 10.1086/713708

Friis, J. I., Sabino, J., Santos, P., Dabelsteen, T., Cardoso, G. C., & Jennions, M. D. (2022). Ecological adaptation and birdsong: how body and bill sizes affect passerine sound frequencies. Behavioral Ecology, 33(4), 798–806. 10.1093/beheco/arac042

Gill, F., Donsker, D., & Rasmussen, P. (2020). *IOC World Bird List* (v 10.2). 10.14344/IOC.ML.10.2.

Gill, F., D. Donsker, and P. Rasmussen. 2023. IOC World Bird List (v 13.1).

Grant, P. R., & Grant, B. R. (2006). Evolution of character displacement in Darwin’s finches. Science, 313, 224–226. 10.1126/science.1128374

Gray, D. A., & Cade, W. H. (2000). Sexual selection and speciation in field crickets. Proceedings of the National Academy of Sciences of the USA, 97, 14449–14454. 10.1073/pnas.97.26.14449

Harvey, M. G., Singhal, S., & Rabosky, D. L. (2019). Beyond reproductive isolation: demographic controls on the speciation process. Annual Review of Ecology, Evolution, and Systematics, 50(1), 75–95. 10.1146/annurev-ecolsys-110218-024701

Hay, E. M., McGee, M. D., & Chown, S. L. (2022). Geographic range size and speciation in honeyeaters. BMC Ecology and Evolution, 22, 1–14. 10.1186/s12862-022-02041-6

Hay, E. M., McGee, M. D., White, C. R., & Chown, S. L. (2024). Body size shapes song in honeyeaters. Proceedings of the Royal Society B: Biological Sciences. 10.1098/rspb.2024.0339

Hernández-Hernández, T., Miller, E. C., Roman-Palacios, C., & Wiens, J. J. (2021). Speciation across the Tree of Life. Biological Reviews, 96(4), 1205–1242. 10.1111/brv.12698

Higgins, P. J., Christidis, L., & Ford, H. (2020a). Little Friarbird (*Philemon citreogularis*), version 1.0. In J. del Hoyo, A. Elliott, J. Sargatal, D. A. Christie, and E. de Juana (Eds.), Birds of the World. Cornell Lab of Ornithology. 10.2173/bow.litfri1.01

Higgins, P. J., Christidis, L., & Ford, H. (2020b). Gray Friarbird (*Philemon kisserensis*), version 1.0. In J. del Hoyo, A. Elliott, J. Sargatal, D. A. Christie, & E. de Juana (Eds.), Birds of the World. Cornell Lab of Ornithology. 10.2173/bow.gryfri1.01

Higgins, P. J., Christidis, L., & Ford, H. (2020c). Yellow-throated Miner (*Manorina flavigula*), version 1.0. In J. del Hoyo, A. Elliott, J. Sargatal, D. A. Christie, & E. de Juana (Eds.). Birds of the World. Cornell Lab of Ornithology. 10.2173/bow.yetmin1.01

Higgins, P. J., Christidis, L., & Ford, H. (2020d). Noisy Miner (*Manorina melanocephala*), version 1.0. In J. del Hoyo, A. Elliott, J. Sargatal, D. A. Christie, & E. de Juana (Eds.). Birds of the World. Cornell Lab of Ornithology. 10.2173/bow.noimin1.01

Higgins, P. J., Christidis, L., & Ford, H. (2020e). Black-eared Miner (*Manorina melanotis*), version 1.0. In J. del Hoyo, A. Elliott, J. Sargatal, D. A. Christie, & E. de Juana (Eds.). Birds of the World. Cornell Lab of Ornithology. 10.2173/bow.blemin1.01

Higgins, P. J., Christidis, L., & Ford, H. (2022). Brown Honeyeater (*Lichmera indistincta*), version 1.1. In S. M. Billerman (Ed.), Birds of the World. Cornell Lab of Ornithology, Ithaca, NY, USA. 10.2173/bow.brohon1.01.1

Higgins, P. J., Ford, H. A., & Christidis, L. (2008). Family Meliphagidae. In J. del Hoyo, A. Elliott, & D. A. Christie (Eds.), Handbook of the Birds of the World: Penduline-tits to shrikes (Vol. 13, pp. 498–691). Lynx Edicions.

Höhna, S., Heath, T. A., Boussau, B., Landis, M. J., Ronquist, F., & Huelsenbeck, J. P. (2014). Probabilistic graphical model representation in phylogenetics. Systematic Biology, 63(5), 753–771. 10.1093/sysbio/syu039

Höhna, S., Landis, M. J., Heath, T. A., Boussau, B., Lartillot, N., Moore, B. R., Huelsenbeck, J. P., & Ronquist, F. (2016). RevBayes: Bayesian phylogenetic inference using graphical models and an interactive model-specification language. Systematic Biology, 65(4), 726–736. 10.1093/sysbio/syw021

Höhna, S., Freyman, W. A., Nolen, Z., Huelsenbeck, J. P., May, M. R., and Moore, B. R. (2019). A Bayesian approach for estimating branch-specific speciation and extinction rates. bioRxiv. 10.1101/555805

Irwin, D. E., Bensch, S., & Price, T. D. (2001). Speciation in a ring. Nature, 409, 333–337. 10.1038/35053059

Jarvis, E. D. (2021). At the beginning of speciation. Science, 371(6536), 1312. 10.1126/science.abg5454

Lachlan, R. F., & Servedio, M. R. (2004). Song learning accelerates allopatric speciation. Evolution, 58(9), 2049–2063. 10.1111/j.0014-3820.2004.tb00489.x

Laidre, M. E., & Johnstone, R. A. (2013). Animal signals. Current Biology, 23(18), R829–833. 10.1016/j.cub.2013.07.070

Maan, M. E., & Seehausen, O. (2011). Ecology, sexual selection and speciation. Ecology Letters, 14(6), 591–602. 10.1111/j.1461-0248.2011.01606.x

Mahler, D. L., Revell, L. J., Glor, R. E., & Losos, J. B. (2010). Ecological opportunity and the rate of morphological evolution in the diversification of Greater Antillean anoles. Evolution, 64(9), 2731–2745. 10.1111/j.1558-5646.2010.01026.x

Marki, P. Z., Jonsson, K. A., Irestedt, M., Nguyen, J. M. T., Rahbek, C., & Fjeldsa, J. (2017). Supermatrix phylogeny and biogeography of the Australasian Meliphagides radiation (Aves: Passeriformes). Molecular Phylogenetics and Evolution, 107, 516–529. 10.1016/j.ympev.2016.12.021

Marquet, P. A., Fernández, M., Navarette, A. A., & Valdovinos, C. (2004). Diversity emerging: towards a deconstruction of biodiversity patterns. In L. M. & H. L. (Eds.), Frontiers of biogeography: New directions in the geography of nature (pp. 191–209). Sinauer Associates.

Marshall, D. C., Slon, K., Cooley, J. R., Hill, K. B. R., & Simon, C. (2008). Steady Plio-Pleistocene diversification and a 2-million-year sympatry threshold in a New Zealand cicada radiation. Molecular Phylogenetics and Evolution, 48(3), 1054–1066. 10.1016/j.ympev.2008.05.007

Mason, N. A., & Burns, K. J. (2015). The effect of habitat and body size on the evolution of vocal displays in Thraupidae (tanagers), the largest family of songbirds. Biological Journal of the Linnean Society, 114, 538–551. 10.1111/bij.12455

Mason, N. A., Burns, K. J., Tobias, J. A., Claramunt, S., Seddon, N., & Derryberry, E. P. (2017). Song evolution, speciation, and vocal learning in passerine birds. Evolution, 71(3), 786–796. 10.1111/evo.13159

Matute, D. R., & Cooper, B. S. (2021). Comparative studies on speciation: 30 years since Coyne and Orr. Evolution, 75(4), 764–778. 10.1111/evo.14181

May, M. R., & Moore, B. R. 2020. A Bayesian approach for inferring the impact of a discrete character on rates of continuous-character evolution in the presence of background-rate variation. Systematic Biology, 69:530–544. 10.1093/sysbio/syz069

Mikula, P., Valcu, M., Brumm, H., Bulla, M., Forstmeier, W., Petruskova, T., Kempenaers, B., & Albrecht, T. (2021). A global analysis of song frequency in passerines provides no support for the acoustic adaptation hypothesis but suggests a role for sexual selection. Ecology Letters, 24(3), 477–486. 10.1111/ele.13662

Mynard, P., Algar, A., Lancaster, L., Bocedi, G., Fahri, F., Gubry-Rangin, C., Lupiyaningdyah, P., Nangoy, M., Osborne, O., Papadopulos, A., Sudiana, I. M., Juliandi, B., Travis, J., & Herrera-Alsina, L. (2023). Impact of phylogenetic tree completeness and misspecification of sampling fractions on trait dependent diversification models. Systematic Biology. 10.1093/sysbio/syad001

Oliveros, C. H., Field, D. J., Ksepka, D. T., Barker, F. K., Aleixo, A., Andersen, M. J., Alstrom, P., Benz, B. W., Braun, E. L., Braun, M. J., Bravo, G. A., Brumfield, R. T., Chesser, R. T., Claramunt, S., Cracraft, J., Cuervo, A. M., Derryberry, E. P., Glenn, T. C., Harvey, M. G., Hosner, P. A., Joseph, L., Kimball, R. T., Mack, A. L., Miskelly, C. M., Peterson, A. T., Robbins, M. B., Sheldon, F. H., Silveira, L. F., Smith, B. T., White, N. D., Moyle, R. G., & Faircloth, B. C. (2019). Earth history and the passerine superradiation. Proceedings of the National Academy of Sciences of the USA, 116(16), 7916–7925. 10.1073/pnas.1813206116

Pagel, M. (1999). Inferring the historical patterns of biological evolution. Nature, 401(6756), 877–884.

Paradis, E., Claude, J., & Strimmer, K. (2004). APE: analyses of phylogenetics and evolution in R language. Bioinformatics, 20(2), 289–290. 10.1093/bioinformatics/btg412

Pearse, W. D., Morales-Castilla, I., James, L. S., Farrell, M., Boivin, F., & Davies, T. J. (2018). Global macroevolution and macroecology of passerine song. Evolution, 72(4), 944–960. 10.1111/evo.13450

Peñalba, J. V., Joseph, L., & Moritz, C. (2019). Current geography masks dynamic history of gene flow during speciation in northern Australian birds. Molecular Ecology, 28, 630–643. 10.1111/mec.14978

Pfennig, D. W., & Pfennig, K. S. (2010). Character displacement and the origins of diversity. The American Naturalist, 176, 26–44. 10.1086/657056

Pfennig, D. W., Wund, M. A., Snell-Rood, E. C., Cruickshank, T., Schlichting, C. D., & Moczek, A. P. (2010). Phenotypic plasticity’s impacts on diversification and speciation. Trends in Ecology & Evolution, 25(8), 459–467. 10.1016/j.tree.2010.05.006

Podos, J. (2001). Correlated evolution of morphology and vocal signal structure in Darwin’s fnches. Nature, 409, 185–188. 10.1038/35051570

Podos, J., & Nowicki, S. (2004). Beaks, adaptation, and vocal evolution in Darwin’s finches. Bioscience, 54, 501–510. 10.1641/0006-3568(2004)054[0501:BAAVEI#x005D;2.0.CO;2

Price, T. (2008). Speciation in Birds. Roberts and Co.

R Core Team. (2013). R: a language and environment for statistical computing. https://www.r-project.org/

Rabosky, D. L., Santini, F., Eastman, J., Smith, S. A., Sidlauskas, B., Chang, J., & Alfaro, M. E. (2013). Rates of speciation and morphological evolution are correlated across the largest vertebrate radiation. Nature Communications, 4, 1958. 10.1038/ncomms2958

Rambaut, A., Drummond, A. J., Xie, D., Baele, G., & Suchard, M. A. (2018). Posterior summarization in Bayesian phylogenetics using Tracer 1.7. Systematic Biology, 67(5), 901–904. 10.1093/sysbio/syy032

Revell, L. J. (2012). phytools: an R package for phylogenetic comparative biology (and other things). Methods in Ecology and Evolution, 3(2), 217–223. 10.1111/j.2041-210X.2011.00169.x

Revell, L. J., Harmon, L. J., & Collar, D. C. (2008). Phylogenetic signal, evolutionary process, and rate. Systematic Biology, 57(4), 591–601. 10.1080/10635150802302427

Ricklefs, R. E. (2006). Global variation in the diversification rate of passerine birds. Ecology, 87(10), 2468–2478. 10.1890/0012-9658(2006)87[2468:GVITDR#x005D;2.0.CO;2

Ritchie, M. G. (2007). Sexual selection and speciation. Annual Review of Ecology, Evolution, and Systematics, 38, 79–102. 10.1146/annurev.ecolsys.38.091206.095733

Schluter, D. 2024. Variable success in linking micro and macroevolution. Evolutionary Journal of the Linnean Society, kzae016. 10.1093/evolinnean/kzae016

Seddon, N., & Tobias, J. A. (2007). Song divergence at the edge of Amazonia: an empirical test of the peripatric speciation model. Biological Journal of the Linnean Society, 90, 173–188. 10.1111/j.1095-8312.2007.00753.x

Servedio, M. R., & Boughman, J. W. (2017). The role of sexual selection in local adaptation and speciation. Annual Review of Ecology, Evolution, and Systematics, 48, 85–109. 10.1146/annurev-ecolsys-110316-022905

Simões, T. R., Vernygora, O., Caldwell, M. W., & Pierce, S. E. (2020). Megaevolutionary dynamics and the timing of evolutionary innovation in reptiles. Nature Communications, 11(1), 3322. 10.1038/s41467-020-17190-9

Singhal, S., Huang, H., Grundler, M. R., Marchan-Rivadeneira, M. R., Holmes, I., Title, P. O., Donnellan, S. C., & Rabosky, D. L. (2018). Does population structure predict the rate of speciation? a comparative test across Australia’s most diverse vertebrate radiation. The American Naturalist, 192(4), 432–447. 10.1086/699515

Suthers, R. A., & Zollinger, S. A. (2008). From brain to song: the vocal organ and vocal tract. In P. H. Zeigler & P. Marler (Eds.), Neuroscience of Birdsong (pp. 78–98). Cambridge University Press.

Tinghitella, R. M., Broder, E. D., Gurule-Small, G. A., Hallagan, C. J., & Wilson, J. D. (2018). Purring crickets: the evolution of a novel sexual signal. The American Naturalist, 192(6), 773–782. 10.1086/700116

Thomas, G. H., and R. P. Freckleton. 2012. MOTMOT: models of trait macroevolution on trees. Methods in Ecology and Evolution, 3:145–151. 10.1111/j.2041-210X.2011.00132.x

Tobias, J. A., Ottenburghs, J., & Pigot, A. L. (2020). Avian diversity: speciation, macroevolution, and ecological function. Annual Review of Ecology, Evolution, and Systematics, 51(1), 533–560. 10.1146/annurev-ecolsys-110218-025023

Tribble, C. M., Freyman, W. A., Landis, M. J., Lim, J. Y., Barido-Sottani, J., Kopperud, B. T., Hӧhna, S., & May, M. R. (2021). RevGadgets: an R package for visualizing Bayesian phylogenetic analyses from RevBayes. Methods in Ecology and Evolution, 13(2), 314–323. 10.1111/2041-210x.13750

Wagner, C. E., Harmon, L. J., & Seehausen, O. (2012). Ecological opportunity and sexual selection together predict adaptive radiation. Nature, 487(7407), 366–369. 10.1038/nature11144

Wilkins, M. R., Seddon, N., & Safran, R. J. (2013). Evolutionary divergence in acoustic signals: causes and consequences. Trends in Ecology & Evolution, 28(3), 156–166. 10.1016/j.tree.2012.10.002

Zuk, M., Rotenberry, J. T., & Tinghitella, R. M. (2006). Silent night: adaptive disappearance of a sexual signal in a parasitized population of field crickets. Biology Letters, 2(4), 521–524. 10.1098/rsbl.2006.0539

