## Supplementary material for "Shifts in diversification influence rates of song evolution in honeyeaters": Figures S1-S7

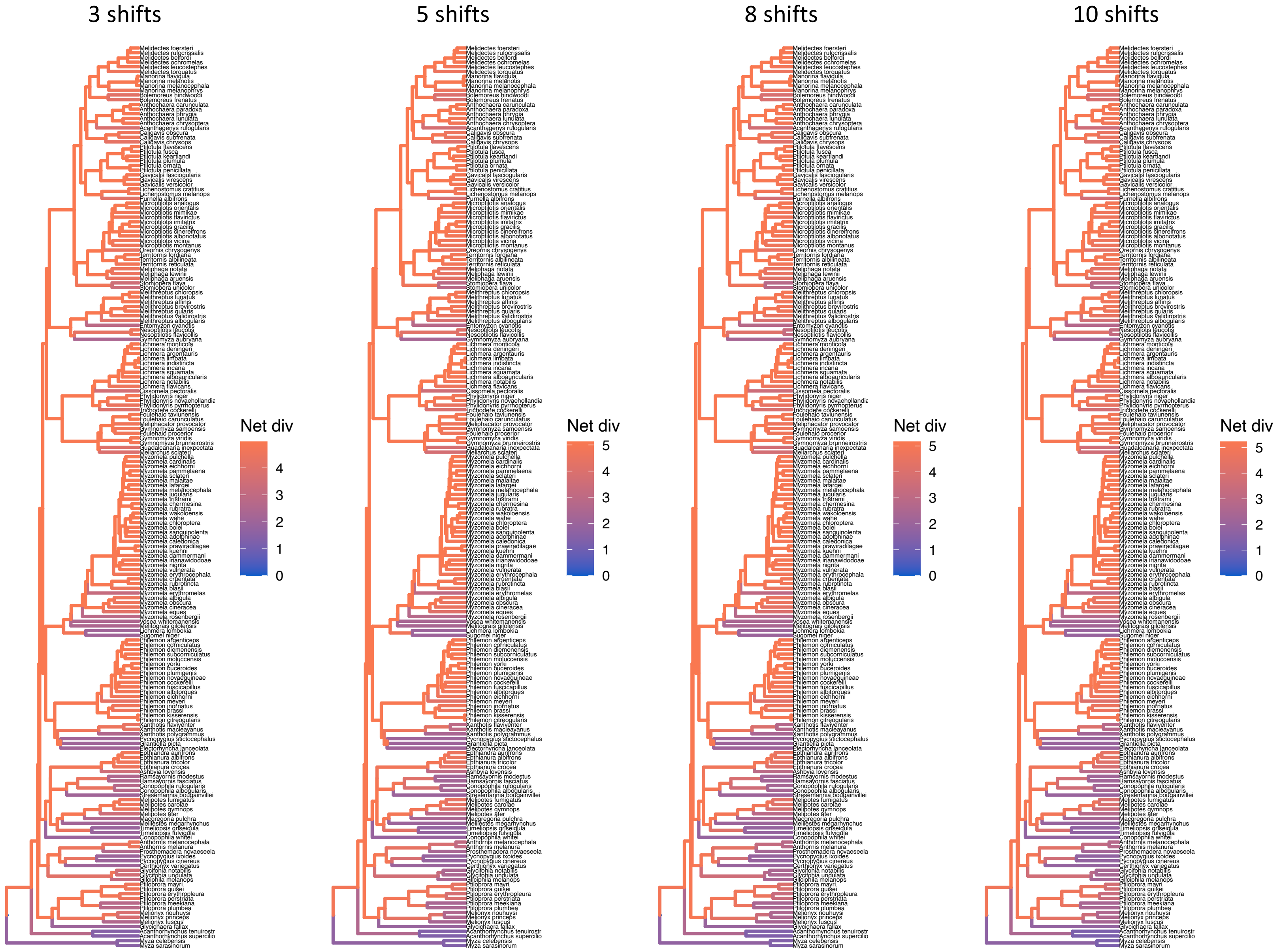


**Figure S1.** Net diversification rate estimates from the branch specific speciation and extinction model in RevBayes, testing different priors. The phylogeny is the full honeyeater tree that contains 192 species.

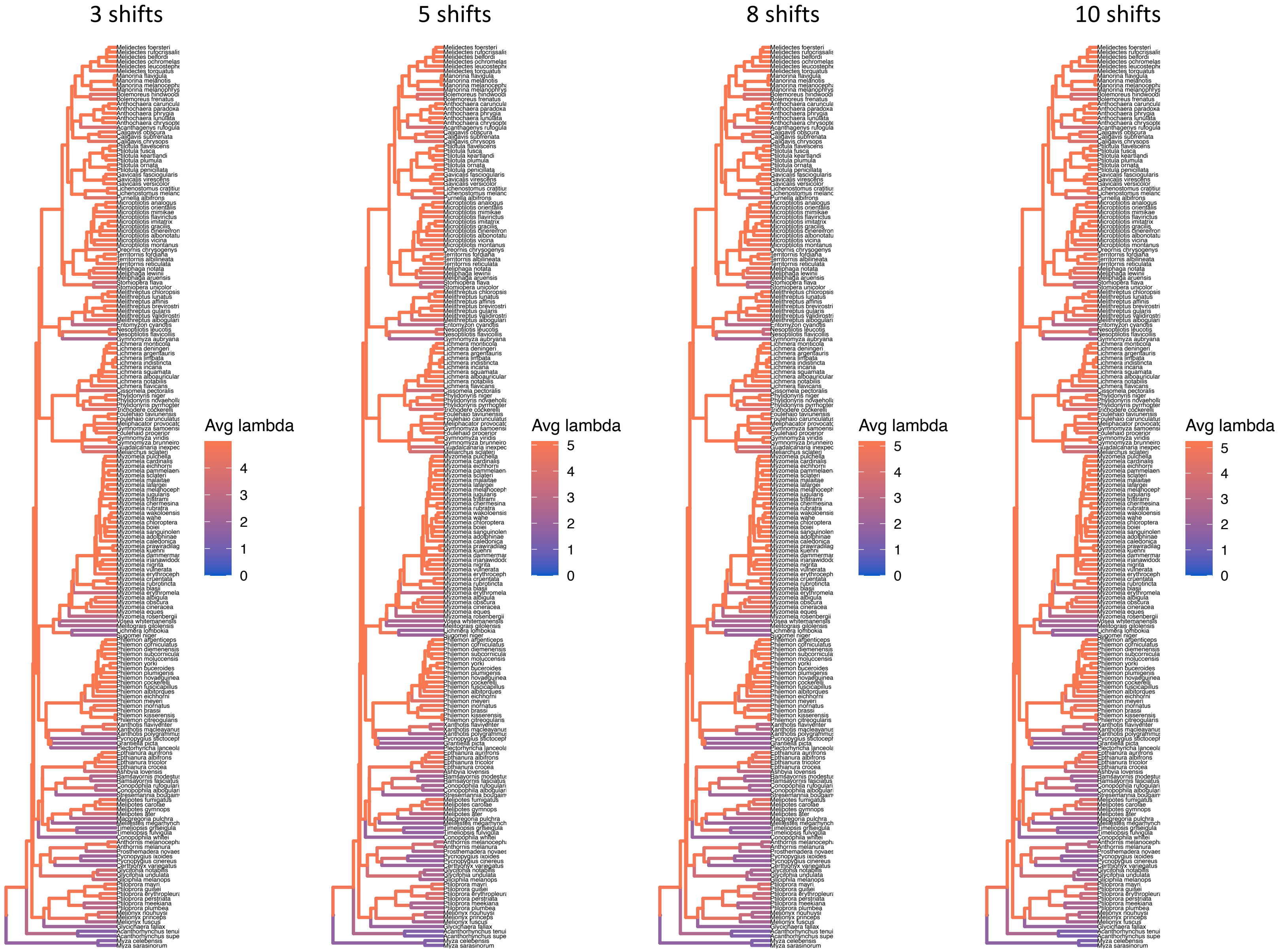


**Figure S2.** Speciation rate estimates from the branch specific speciation and extinction model in RevBayes, testing different priors. The phylogeny is the full honeyeater tree that contains 192 species.

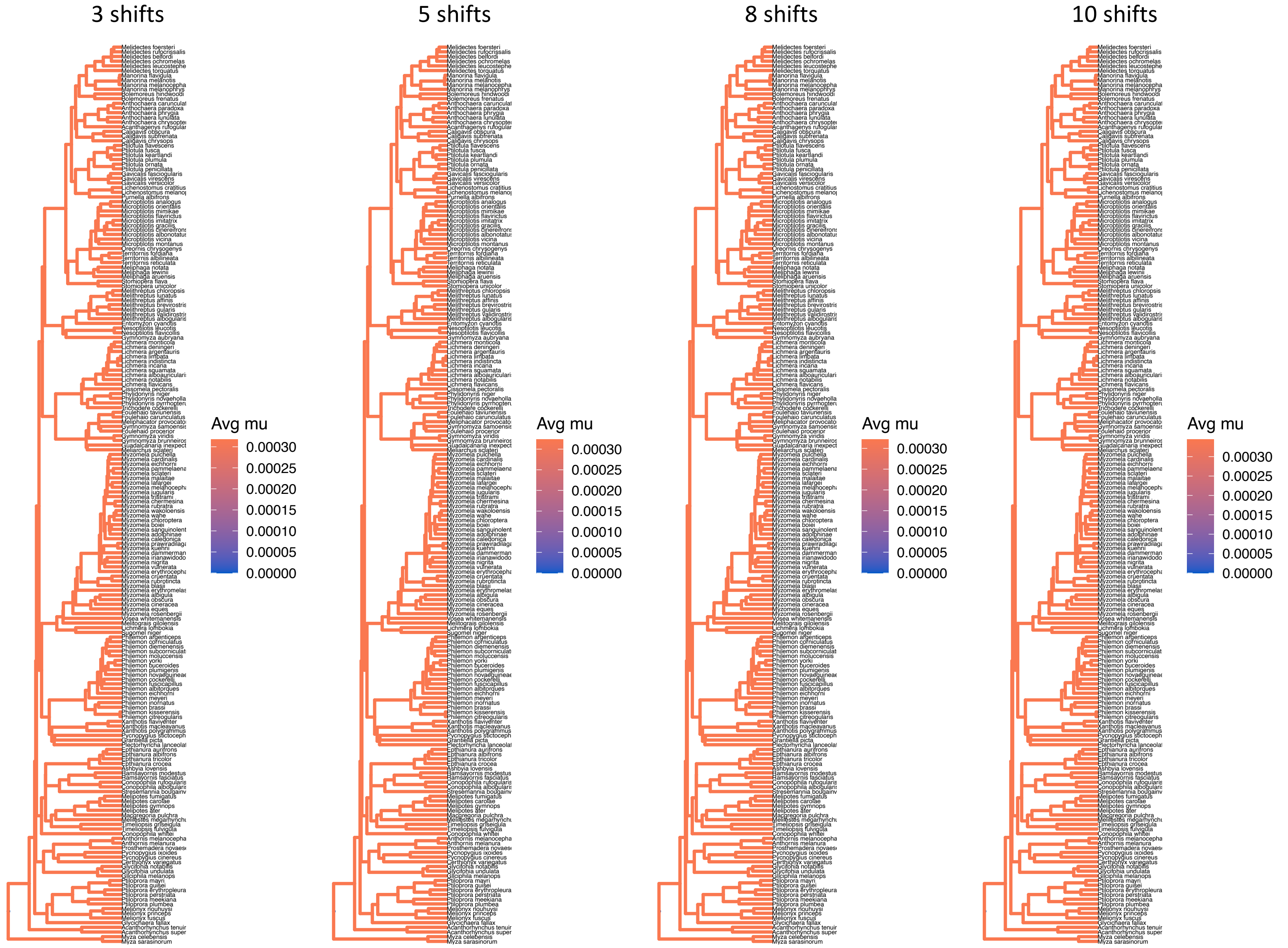


**Figure S3.** Extinction rate estimates from the branch specific speciation and extinction model in RevBayes, testing different priors. The phylogeny is the full honeyeater tree that contains 192 species.

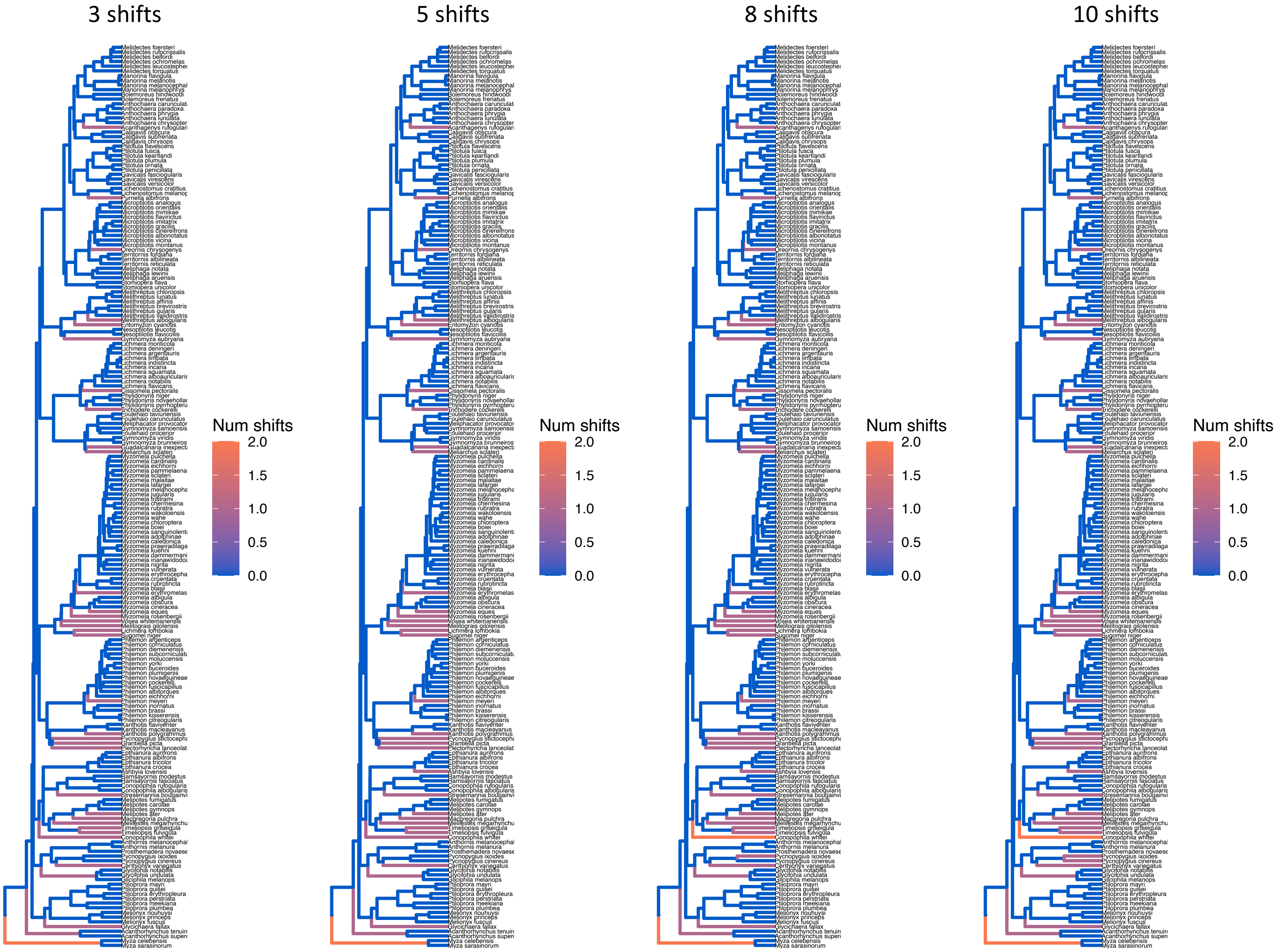


**Figure S4.** Shifts in diversification from the branch specific speciation and extinction model in RevBayes, testing different priors. The phylogeny is the full honeyeater tree that contains 192 species.

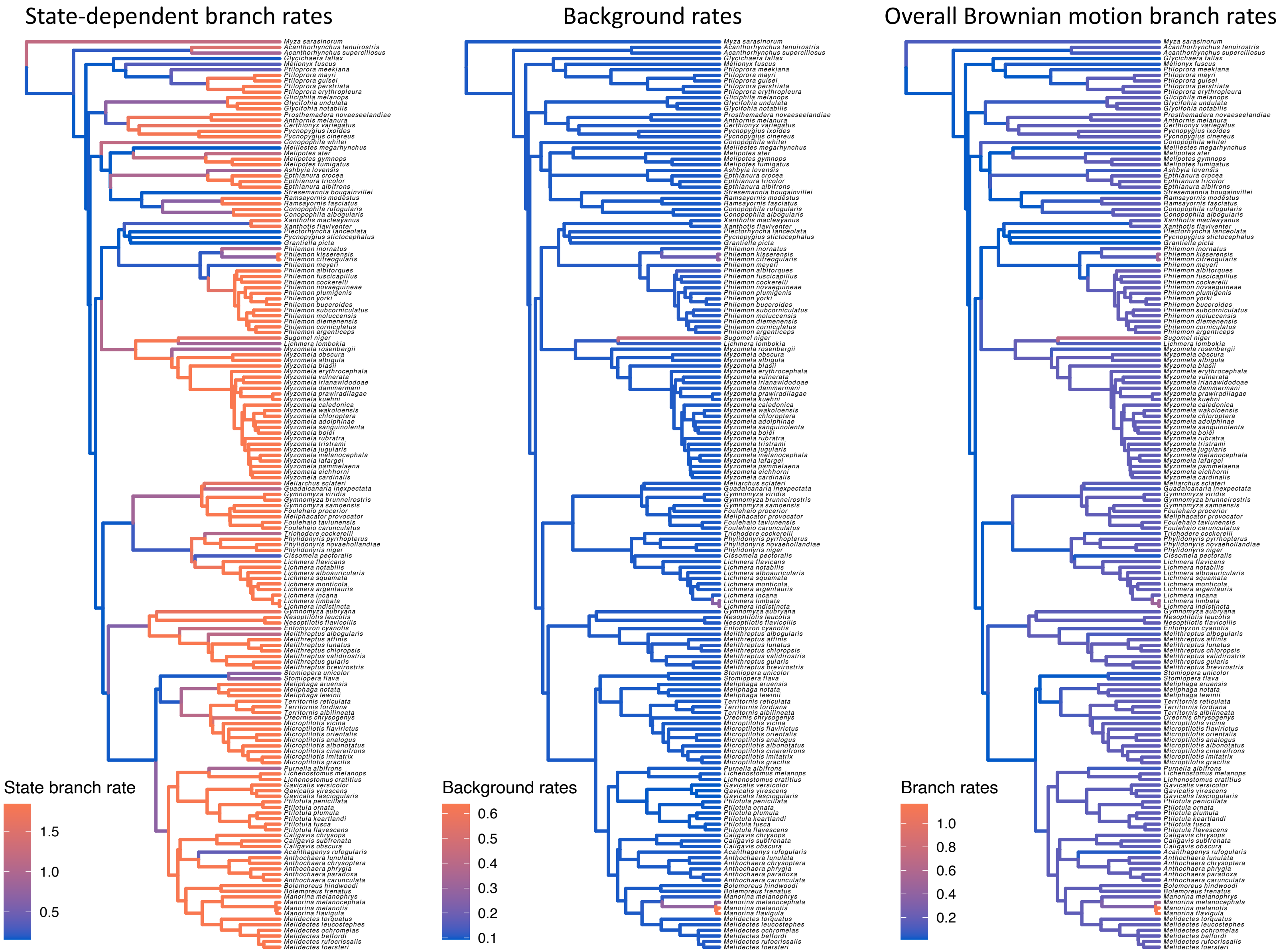


**Figure S5.** Estimated Brownian-motion rates of song evolution with state-dependent and background-rate variation for 163 species of honeyeaters. This model had a prior of 5 rate shifts for the evolution of the continuous characters. Left) displays state-dependent rates of continuous-character evolution. Middle) Background rates of continuous-character evolution, highlighting rate variation that cannot be attributed to the state of the discrete character. Right) We show the combined effect of state-dependent and state-independent sources of rate variation, analogous to a Brownian motion model of trait evolution on the song characters.

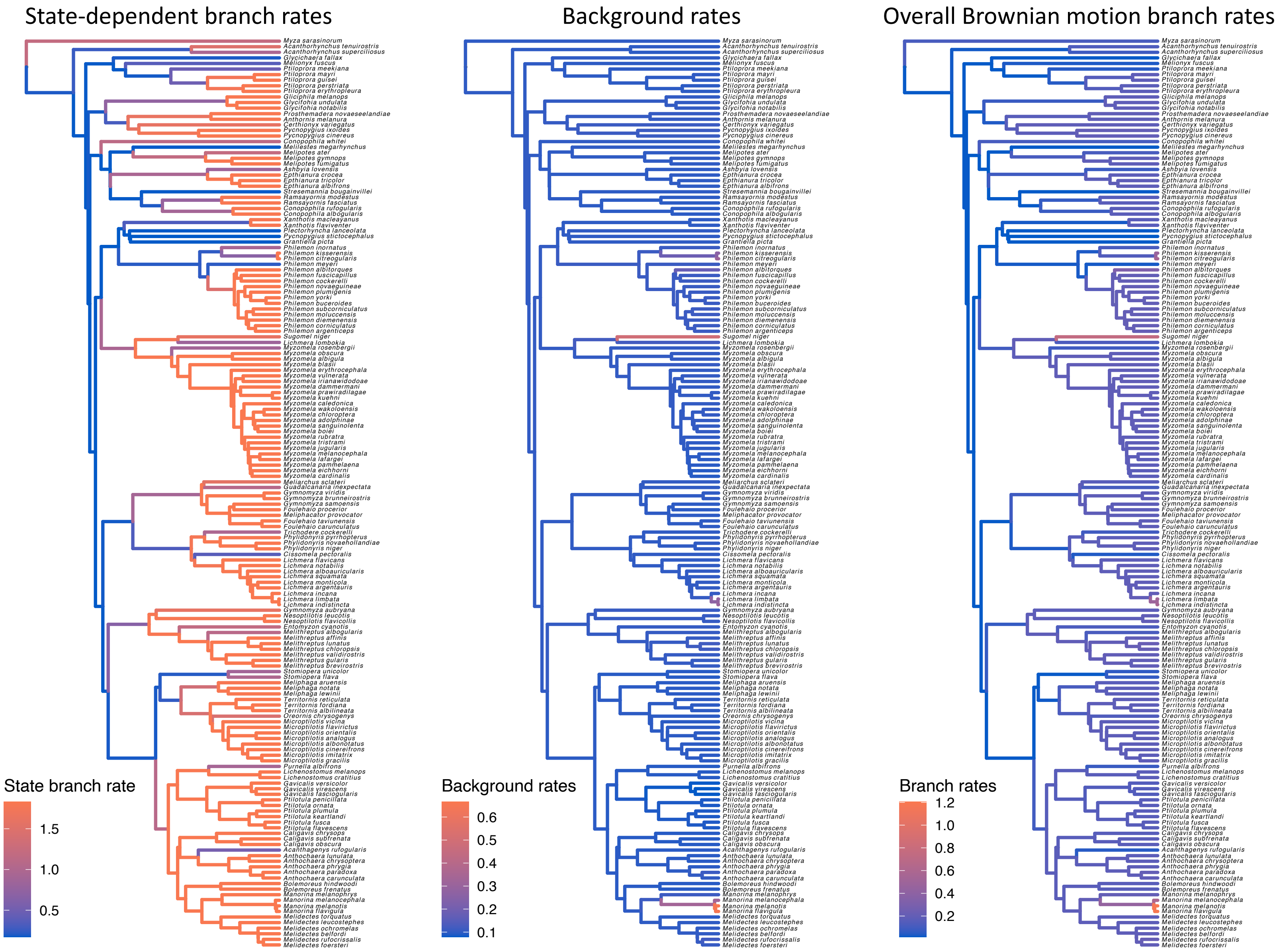


**Figure S6.** Estimated Brownian-motion rates of song evolution with state-dependent and background-rate variation for 163 species of honeyeaters. This model had a prior of 10 rate shifts for the evolution of the continuous characters. Left) displays state-dependent rates of continuous-character evolution. Middle) Background rates of continuous-character evolution, highlighting rate variation that cannot be attributed to the state of the discrete character. Right) We show the combined effect of state-dependent and state-independent sources of rate variation, analogous to a Brownian motion model of trait evolution on the song characters.

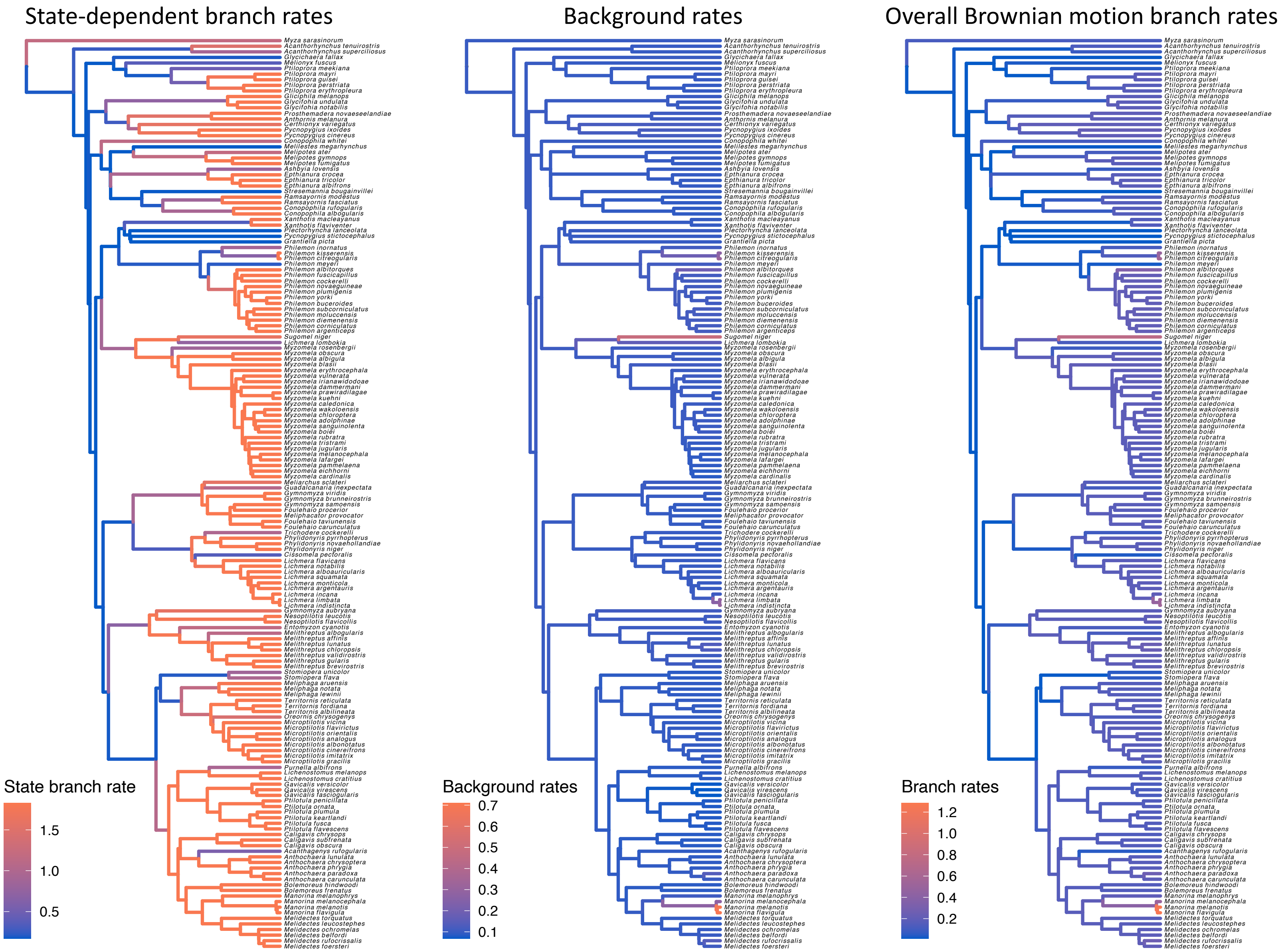


**Figure S7.** Estimated Brownian-motion rates of song evolution with state-dependent and background-rate variation for 163 species of honeyeaters. This model had a prior of 15 rate shifts for the evolution of the continuous characters. Left) displays state-dependent rates of continuous-character evolution. Middle) Background rates of continuous-character evolution, highlighting rate variation that cannot be attributed to the state of the discrete character. Right) We show the combined effect of state-dependent and state-independent sources of rate variation, analogous to a Brownian motion model of trait evolution on the song characters.
